# Modelling the effect of V1a receptor antagonism and its potential therapeutic effect in circadian disorders

**DOI:** 10.1101/2025.11.07.687173

**Authors:** Marcelo Boareto, Jorge Mendoza, Sebastian C. Holst, Andries Kalsbeek, Eric Prinssen, Christophe Grundschober

## Abstract

**Background:** The suprachiasmatic nucleus (SCN) is the central circadian clock in mammals, regulating many daily physiological and behavioral rhythms. Dysregulation of the SCN is associated with various circadian disorders, highlighting the potential therapeutic benefits of targeting its neurons and output pathways. Vasopressin signaling is one of the main regulators of the synchronicity and the functional output of the SCN.

**Methods:** We investigated the effect of a single dose (30mg/kg) of the vasopressin V1a receptor (V1aR) antagonist, balovaptan, on resynchronization of locomotor activity rhythms in mice after a 6-hour phase advance of the light-dark cycle. To mechanistically model the effect of V1aR antagonism, we developed a mathematical framework simulating the SCN, its control of circadian biomarkers (melatonin, core body temperature), and the impact of V1aR antagonism.

**Results:** A single administration of the V1a antagonist balovaptan significantly accelerated resynchronization of locomotor activity rhythms to new light-dark cycles. To mechanistically understand this effect, we devised a mathematical model of the SCN that successfully captures this accelerated synchronization of circadian rhythms under V1aR antagonism. Additionally, the model replicates well-established SCN behaviors in both humans and rodents, including the phase response curve triggered by a light pulse at various circadian phases, and the desynchronization of the SCN observed in forced desynchronization experiments.

Mechanistically, our model suggests that weakening vasopressin signaling via V1aR antagonism strengthens the SCN’s resistance to internal desynchronization. Additionally, our model suggests a strong link between the endogenous period (tau) and the phase of circadian biomarkers, with longer tau values resulting in delayed biomarker rhythms. Importantly, the model predicts that V1aR antagonism induces a phase advance proportional to tau. The model predicts that individuals with longer endogenous periods, who consequently exhibit greater phase delays in their circadian rhythms, could experience more substantial phase advances in response to V1aR antagonism.

**Discussion:** We show that targeting V1aR is enough to cause a faster resynchronization to a new light-dark cycle in the jet lag paradigm and establish a computational framework for investigating its therapeutic potential in circadian rhythm disorders. This framework, adaptable to incorporate pharmacodynamic data, can be used to design clinical trials evaluating V1aR antagonism for treating circadian disorders.

## Introduction

Circadian rhythms are essential for regulating many physiological processes, including sleep-wake cycles, hormone secretion, and metabolism (Takahashi, 2017). Disruptions in circadian rhythms are associated with numerous health issues, such as sleep disruption, metabolic syndrome, and mood disorders (Czeisler & Gooley, 2007). Understanding the mechanisms underlying circadian regulation is crucial for developing therapeutic strategies to address these conditions.

The suprachiasmatic nucleus (SCN) is located within the anterior hypothalamus and serves as the primary circadian pacemaker in mammals, coordinating daily physiological and behavioral rhythms. This critical structure displays spatial heterogeneity, characterized by distinct neuronal populations in its ventrolateral (core) and dorsomedial (shell) regions. The core region is predominantly populated by vasoactive intestinal peptide (VIP) neurons, which receive direct photic input from intrinsically photoreceptive retinal ganglion cells (ipRGC) via the retinohypothalamic tract, enabling synchronization to environmental light cues. These VIP neurons play a pivotal role in intercellular communication, synchronizing other SCN neurons (Aton et al, 2005, Aton and Herzog, 2005). In contrast, the shell region is enriched with arginine vasopressin (AVP) neurons, which contribute significantly to the rhythmic output of the SCN. The differential distribution of VIP and AVP neurons, coupled with their specific functions, underlies the SCN’s complex ability to regulate circadian rhythms (Hastings, et al, 2003). Moreover, recent single-cell RNA sequencing studies show that SCN heterogeneity is more complex than this core and shell subdivision (Xu et al., 2021; Morris et al., 2021).

Experiments employing non-24-hour light cycles, such as continuous light (Aschoff et al, 1967) or 28-hour light-dark cycles (Dijk & Czeisler, 1995), demonstrate the vulnerability of the SCN to desynchronization. Notably, studies in rodents using 22-hour light-dark cycles reveal a functional dissociation between ventrolateral (vlSCN) VIP and dorsomedial (dmSCN) AVP neurons (de la Iglesia et al., 2004). VIP neurons readily entrain to the 22-hour light cycle, whereas AVP neurons maintain a near-24-hour rhythm, indicating a higher sensitivity of VIP neurons to light cues (de la Iglesia et al., 2004). This diverging entrainment suggests that although VIP neurons can influence AVP neurons, significant deviations between these two populations can lead to desynchronization within the SCN. Consequently, the SCN output is also distinctly affected. For instance, core body temperature, mostly influenced by dmSCN AVP output, tends to follow the endogenous circadian period (Aschoff., 1965; Dijk and Czeisler, 1995; Cambras, et al 2007). Conversely, melatonin, influenced by both vlSCN VIP and dmSCN AVP neurons, exhibits a complex pattern that reflects a combined interplay between these neuronal populations in melatonin regulation (Schwartz et al., 2009).

The strength of arginine vasopressin (AVP) signaling, mediated by V1a and V1b receptors, plays a critical role in synchronizing the suprachiasmatic nucleus (SCN) and entraining circadian rhythms. Notably, co-treatment with V1b and V1aR antagonism, which inhibits AVP signaling, or a dual V1a and V1b knock-out has been shown to accelerate entrainment in mice, particularly in experimental jet lag scenarios (Yamaguchi et al., 2013). This effect has since been reported to be mediated by two independent and additive pathways: an intra-SCN V1a pathway and an extra-SCN V1b pathway (Yamaguchi et al, 2023). This suggests that manipulating AVP signaling could offer a novel approach to mitigating circadian rhythm disruptions.

In this study, we show that acute inhibition of only the V1a vasopressin receptor is sufficient to accelerate circadian resynchronization after a 6 hours phase advance of the light-dark (LD) cycle. Subsequently, we utilized an *in silico* model of the SCN to simulate the impact of V1aR antagonism on key circadian biomarkers, such as melatonin and core body temperature. By integrating experimental data with theoretical modeling, we aim to elucidate the SCN’s response to pharmacological interventions and explore the therapeutic potential of V1aR antagonism for circadian rhythm disorders.

## Results

### Behavioural re-synchronization to a 6h phase-advance of the LD cycle after balovaptan administration

We assessed whether the specific vasopressin V1a antagonist balovaptan (Schnider et al, 2020) might affect light synchronization in animals exposed to an experimental jet lag protocol. Mice (sham, vehicle and V1a antagonist) exposed to a 6-h phase-advance of the light-dark cycle were treated with a single dose of vehicle control administration or 30 mg/kg of the V1a antagonist by oral administration at ZT6 (6h after lights on) on the last day before the shifted LD cycle (Figure 1A).

**Figure 1:**
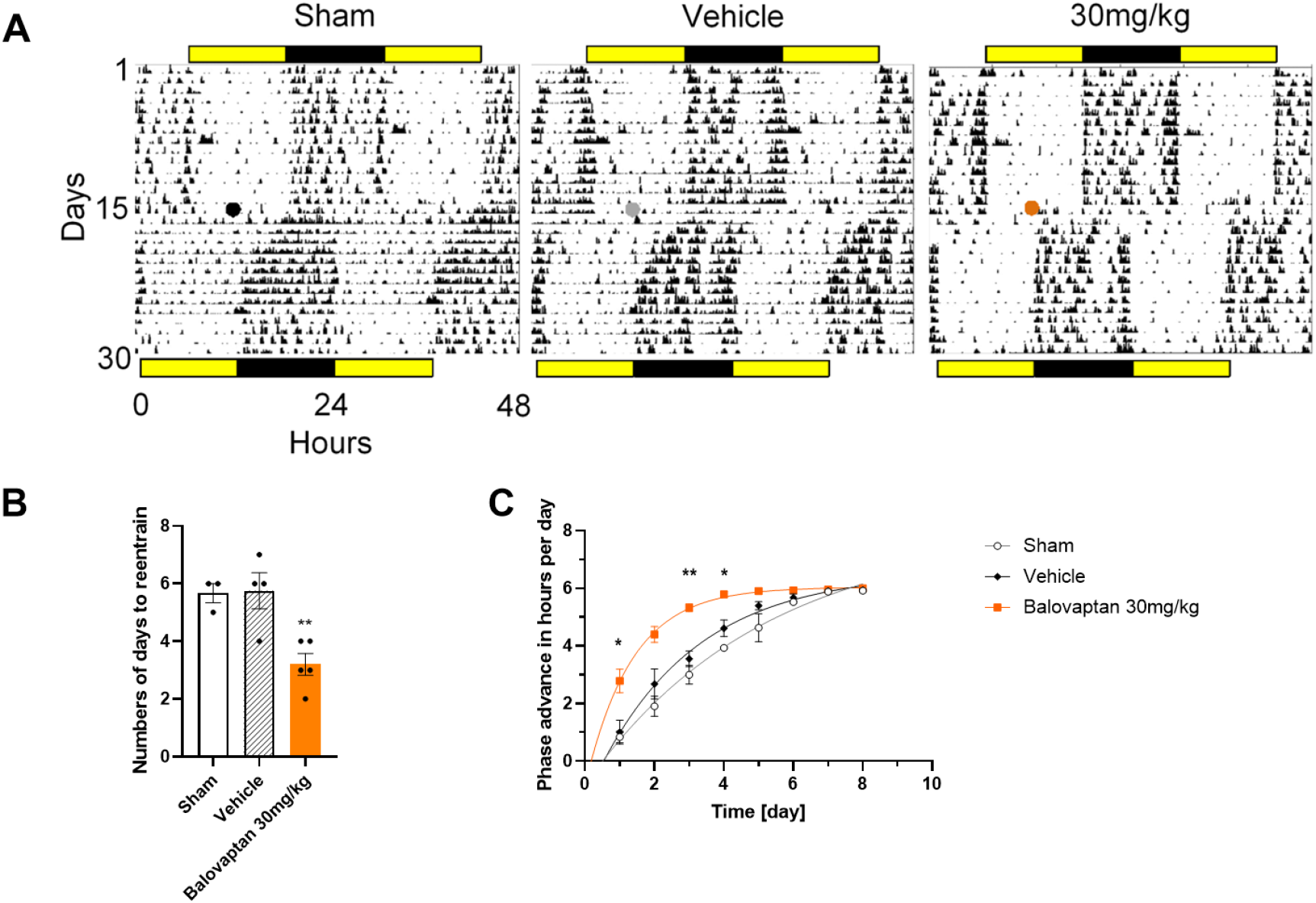
A) Representative actograms of the locomotor activity of mice from each group treated at ZT6 (mid rest period) and exposed to a 6h phase shift of the LD cycle. Color dots in actograms represent time of manipulation, and yellow and black bars in top and bottom indicate the light and dark period, respectively. B) Rate of re-entrainment (number of days) to show full synchronization to the new LD cycle (Dunnett’s post-hoc test vs vehicle.** p= 0.005). C) The speed of entrainment was evaluated daily and on days 1, 3 and 4 the 30 mg/kg group showed a significantly faster re-entrainment than the vehicle group (Dunnett’s post-hoc test vs vehicle.* p< 0.05, ** p< 0.01) (sham n = 3, vehicle n = 4, 30 mg/kg n = 5).

Animals from the control sham group (i.e. only handling for oral administration) needed 5.66 ± 0.33 days to entrain to the new LD cycle. Animals receiving the vehicle administration took 5.75 ± 0.62 days to re-entrain to the new LD cycle. In contrast, mice that were treated with 30 mg/kg balovaptan took only 3.2 ± 0.37 days to show a total behavioural re-entrainment after the 6h phase shift. This effect was statistically significant (F_2,9_ = 10.1, p = 0.005) (Figure 1B). The onset of activity showed a much more rapid daily adjustment to the new LD cycle in 3 of the first 4 days after the shift in the balovaptan treated group (F_2,9_=25, p=0.0002) (Figure 1C). Locomotor activity induced by manipulations was not significantly different between groups (F_2,9_=1.25, p=0.32. Fig S1A supplementary material).

To confirm this finding the experiment was repeated in a new set of animals. The control sham group took 5.33 ± 0.88 days to entrain to the new LD cycle. Animals receiving the vehicle administration took 5.50 ± 0.64 days to re-entrain to the new LD cycle. Similarly to the first jet-lag experiment, mice treated with 30 mg/kg of balovaptan took only 3.40 ± 0.24 days to show a total behavioural reentrainment after the 6h phase shift. This effect was statistically different (F_2,9_=4.99, p = 0.03; Fig. S2A). The onset of activity showed a much more rapid daily adjustment to the LD cycle for the first 3 days after the shift in the V1a antagonist treated group (F_2,9_=14, p= 0.001; Fig. S2B). The three experimental manipulations (sham, vehicle and compound administrations) induced an increase in locomotion, that was not significantly different between groups (F_2,9_=2.91, p=0.10; Fig S1B supplementary material).

### In silico model of V1aR antagonism effect on the SCN

In order to mechanistically understand the impact of V1aR antagonism, we developed a mathematical model to simulate the suprachiasmatic nucleus (SCN), its regulation of key circadian biomarkers (melatonin and core body temperature), and the effect of V1aR antagonism.

At the molecular level, we assume that each neuron has a molecular clock with an endogenous period close to 24 hours, which adjusts its phase in response to various input signals. The model simplifies the molecular clock by representing the overall phase in time without detailing the transcriptional feedback of clock genes. At the cellular level, the model differentiates between VIP and AVP neurons. VIP neurons receive light input and release VIP as neurotransmitter, while AVP neurons receive no light input and release AVP as neurotransmitter. Both neuron types receive inputs from each other, but only VIP neurons are directly influenced by light. These interactions modulate the phase of the molecular clock and regulate the expression of VIP and AVP, which act as synchronization agents. At the tissue level, the SCN is represented as two clusters of cells corresponding to VIP and AVP neurons, reflecting their spatial segregation into core/ventrolateral and shell/dorsomedial regions. These clusters communicate through their respective neurotransmitters, independently modulating the expression of circadian biomarkers and resulting in distinct outcomes for the SCN. The core body temperature is assumed to be controlled only by the output of the shell/dorsomedial region (AVP output), while melatonin is assumed to be controlled by light input and both core/ventrolateral (VIP output) and shell/dorsomedial (AVP output) regions (Figure 2).

**Figure 2.**
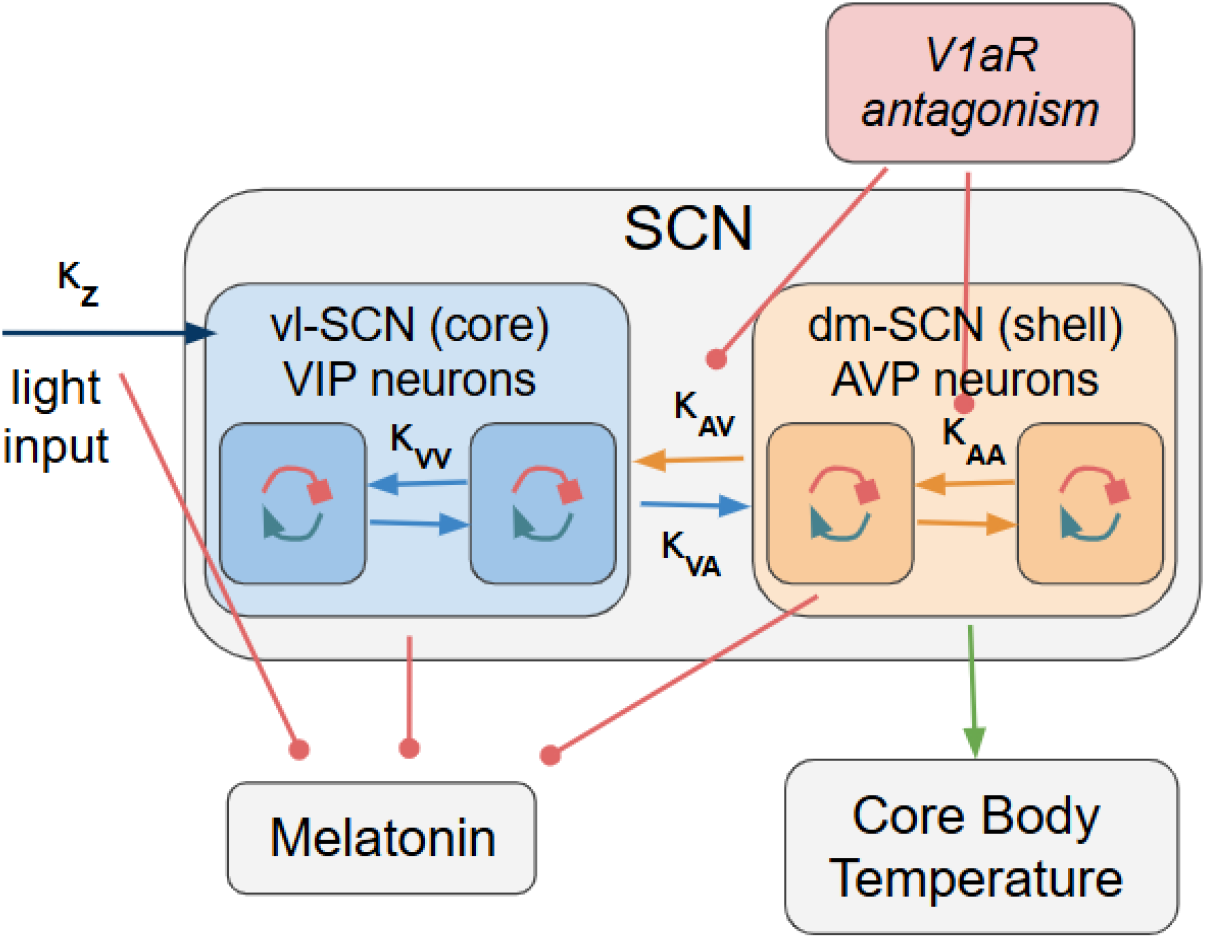
Graphical representation of the suprachiasmatic nucleus (SCN) model, its modulation of circadian biomarkers, and effect of V1aR antagonism. The model represents the SCN as two interacting neuronal populations: ventrolateral (vlSCN - core) VIP and dorsomedial (dmSCN - shell) AVP neurons. These populations communicate via VIP and AVP neurotransmission, respectively, and independently modulate circadian biomarker expression. Core body temperature is exclusively regulated by AVP output, while melatonin is influenced by light input and both VIP and AVP outputs. V1aR antagonism is modeled as a reduction in AVP signaling efficacy. Key parameters: κ_Z_ (light input strength); κ_VV_ (strength of VIP self-modulation); κ_VA_ (strength of VIP modulation of AVP neurons); κ_AA_ (strength of AVP self-modulation); κ_AV_ (strength of AVP modulation of VIP neurons).

The model framework is based on the Kuramoto model (Kuramoto, 1975) describing the temporal evolution of individual cell phases, and many previous models have been a source of inspiration (Amdaoud et al, 2007; Rougemont and Naef 2008; Lu et al, 2016; Kori et al, 2017; Rohling and Meylahn, 2020). For more details and mathematical representation of the model please refer to the Methods section.

### Entrainment to light input (Zeitgeber)

As a first step to validate our mathematical model and translate preclinical findings to humans, we simulated the behavior of SCN under various light conditions to replicate experimental observations and assess the model’s ability to capture the fundamental properties of circadian rhythm regulation.

Under constant dark conditions (DD), where there is no external light input, both VIP and AVP cells were entrained to their intrinsic molecular clocks. The outputs of both cell types overlapped, exhibiting a free-running period of approximately 24.2 hours, consistent with the average endogenous period of the cells (Figure 3A) as defined to match the tau observed in humans. In light dark (LD) conditions, simulating a normal day-night cycle, both VIP and AVP cells in the model entrained to the light input, exhibiting a 24-hour period. This demonstrates the model’s ability to capture the entrainment of circadian rhythms to external light cues, a fundamental feature of the SCN (Figure 3B). Under constant light (LL) conditions, the model showed a break in synchrony between VIP and AVP cells. VIP cells, which receive direct light input, exhibited a longer period, while AVP cells maintained a free-running period close to the endogenous period. Melatonin expression becomes suppressed by both the desynchrony of VIP and AVP output and constant light input, while core body temperature oscillates with a period close to the AVP output and endogenous period (Figure 3C).

**Figure 3.**
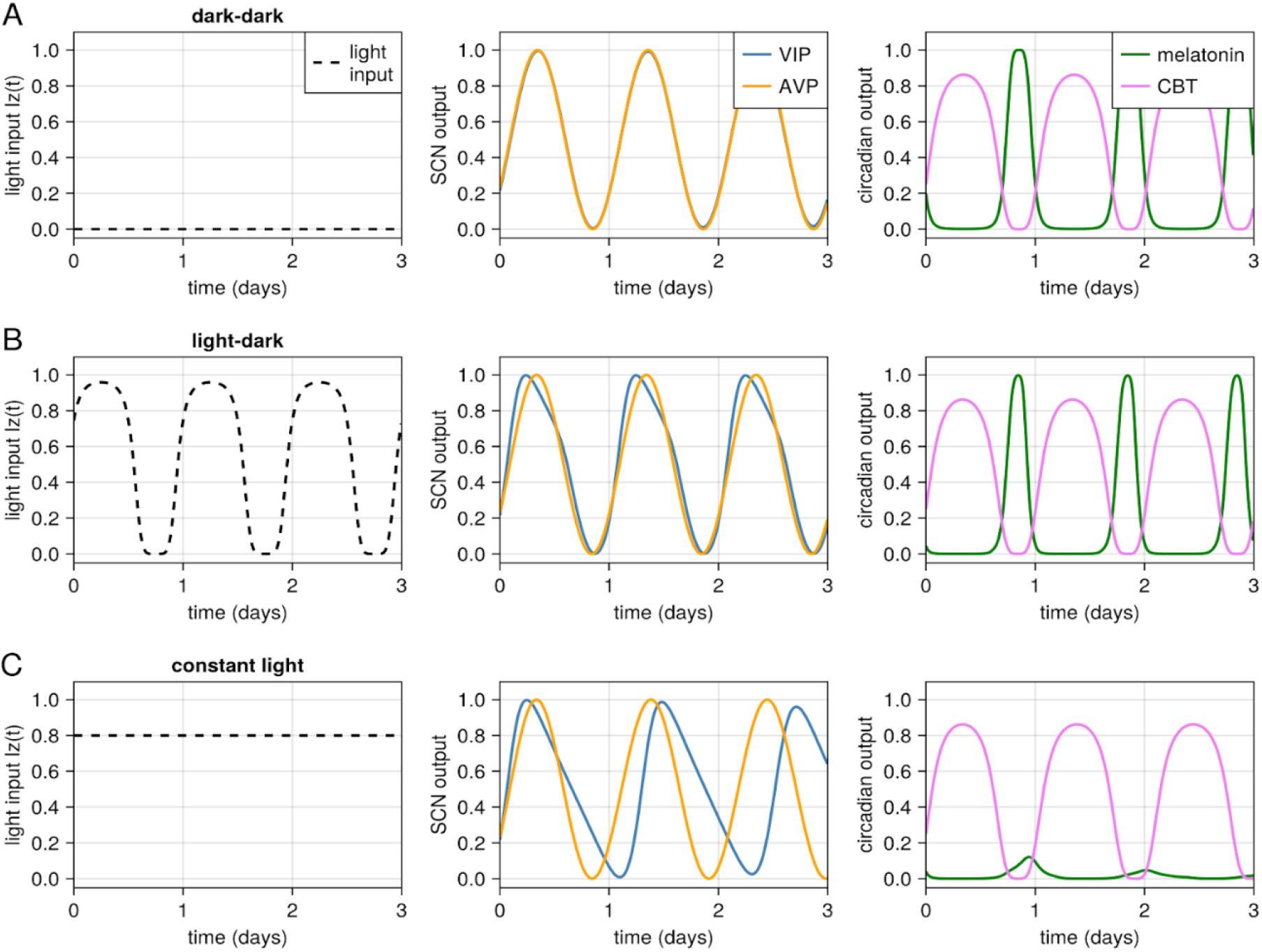
SCN entrainment to varying light conditions. A) Absence of light input (DD). VIP and AVP neurons exhibit a free-running period of approximately 24.2 hours, consistent with their intrinsic molecular clocks, and maintain synchrony. B) Light-dark (LD) cycle. Both VIP and AVP neurons entrain to the 24-hour LD cycle, with VIP neurons slightly phase-advanced relative to AVP neurons. C) Constant light (LL). VIP neurons exhibit a longer period due to direct light input, while AVP neurons maintain a free-running period, leading to desynchronization. Melatonin expression is suppressed by both VIP-AVP desynchrony and light, whereas core body temperature oscillates with a period close to the AVP output and the endogenous period.

### Light pulse and phase response curve

To further validate our model against experimental observations, we simulated the effect of a light pulse administered at different phases of the circadian cycle. This generates a phase response curve (PRC), which illustrates the phase shifts in circadian rhythms induced by a zeitgeber (time cue) applied at various circadian phases. A light pulse administered at different circadian phases elicits distinct effects on the circadian system. Specifically, a light pulse delivered before the minimum of core body temperature (CBT), which serves as the reference point for circadian phase (CP = 0), typically induces a phase delay in circadian rhythms (Khalsa et al., 2003). Conversely, a light pulse administered after CP = 0 results in a phase advance (Khalsa et al., 2003). Our model shows consistency with these experimental observations (Figure 4).

**Figure 4.**
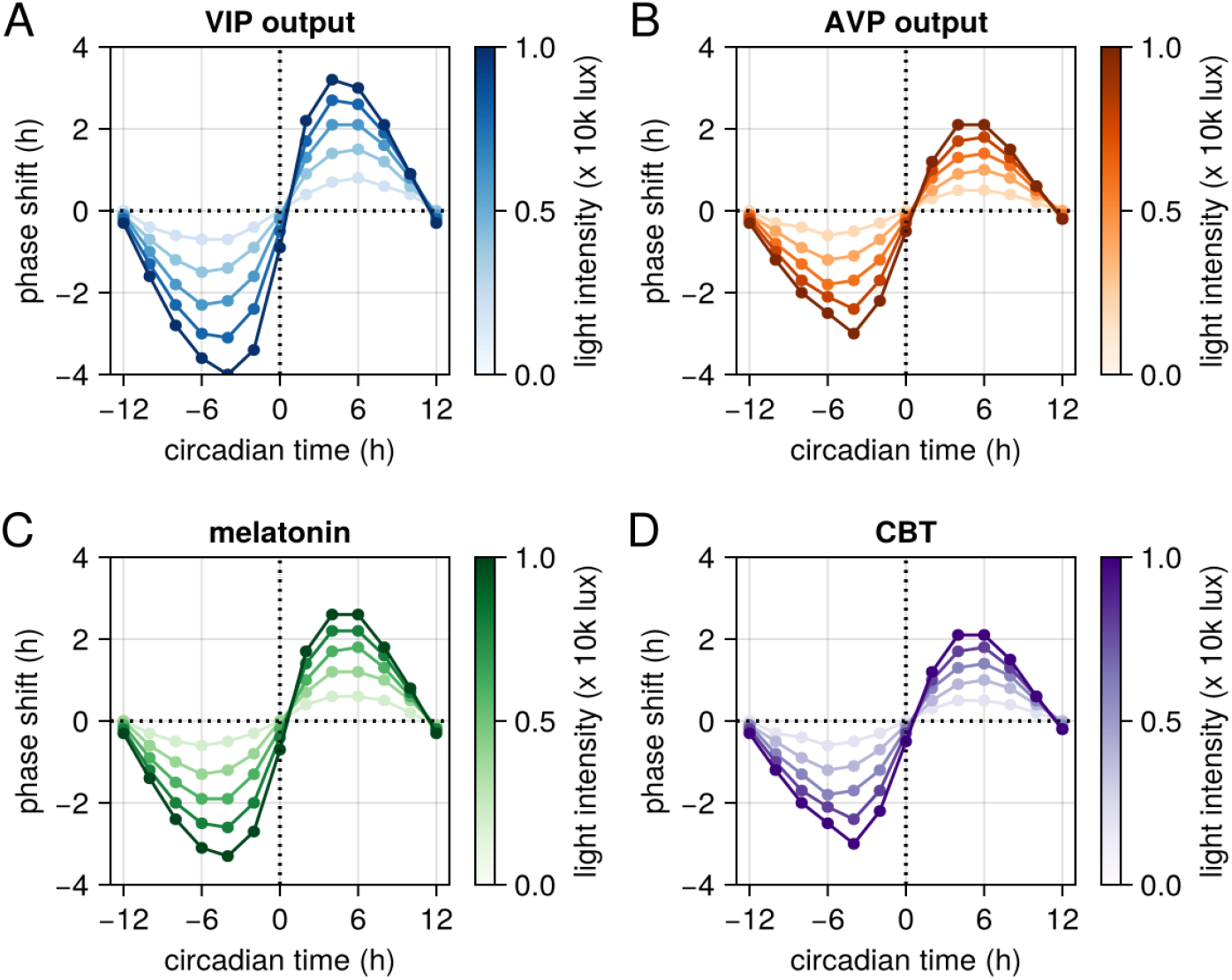
Model-generated phase response curves (PRCs) for SCN neurons and circadian biomarkers. The PRCs illustrate the phase shifts after 3 days induced by a light pulse administered at different circadian phases. The color gradient from light to dark represents the predicted PRC for varying light intensities. PRC for A) VIP neurons, B) AVP neurons, C) melatonin, and D) core body temperature.

Our model also replicates the experimental observation that a light pulse administered before the rise in melatonin levels suppresses melatonin onset and results in a shorter subsequent melatonin peak (Khalsa et al., 2003, Figure S3). This compression of the melatonin profile arises from the desynchronization of VIP and AVP neuronal outputs, both of which regulate melatonin expression. This phenomenon has also been observed in rodents subjected to forced desynchronization experiments (Schwartz et al., 2009).

### Targeting V1aR leads to fast entrainment under jet lag paradigm

As a final step towards validating our model against experimental evidence, we simulated a jet lag paradigm and examined the effects of V1aR antagonism on the rate of entrainment (Figure 5). The model was able to reproduce the accelerated resynchronization of circadian rhythms under V1aR antagonism (Yamagushi et al, 2013; Figure 1). Notably, the model predicts faster synchronization of AVP neuronal output in the presence of V1aR antagonism and no effect of AVP antagonism on the synchronization speed of VIP neurons. This differential response leads to a transient desynchronization between VIP and AVP outputs following the simulated jet lag, consistent with experimental observations in rodents (Davidson, et al 2009; Albus et al, 2005), and previous SCN modelling efforts (Aleixo et al, 2025).

**Figure 5.**
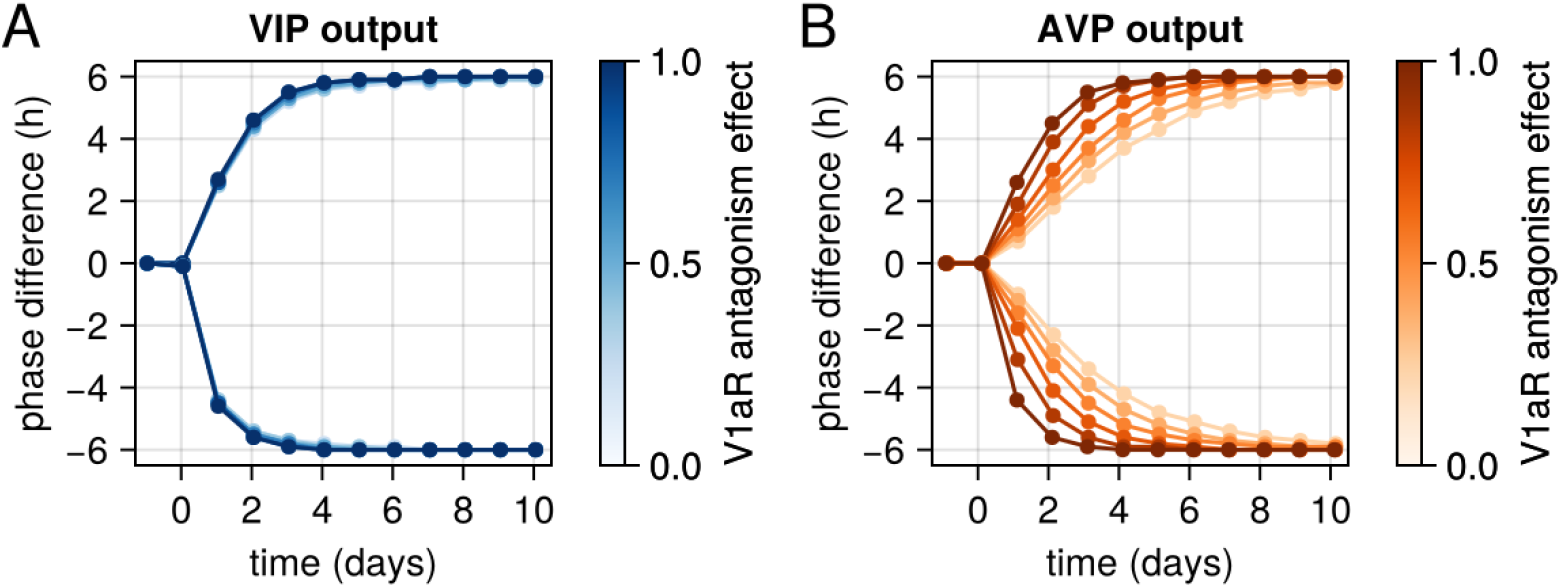
Effects of V1aR antagonism on VIP and AVP neuronal synchronization following simulated jet lag. This figure illustrates the phase differences of VIP and AVP neurons relative to their original schedule after a 6-hour phase advance or delay simulating a jet lag paradigm. A) Phase difference of VIP neurons. B) Phase difference of AVP neurons. The model predicts that without V1aR blockade, VIP neurons recover faster than AVP neurons. However, complete V1aR antagonism eliminates this difference, resulting in a coordinated and rapid resynchronization of both neuronal populations.

### Impact of AVP signaling strength on SCN synchronization

The degree of synchronization between AVP and VIP neurons within the SCN is strongly influenced by both the period of the zeitgeber (external time cue) and the intrinsic endogenous period (tau) of the neurons. Our model shows that AVP and VIP neurons remain synchronized when the zeitgeber period approximates the natural light-dark cycle (∼24 hours). However, when the zeitgeber period deviates significantly from 24 hours (e.g., 22 or 28 hours), desynchronization arises between AVP and VIP neurons (Figure 6A). This desynchronization stems from the differential responses of these neuronal populations: VIP neurons, which receive direct light input, adjust their oscillation period to match the zeitgeber, whereas AVP output tends to be close to the endogenous period (Figure 6A). These results are in agreement with experimental observations in rodents and humans (Dijk & Czeisler, 1995; de la Iglesia et al., 2004), and previous modelling observations (Goltsev et al, 2022).

**Figure 6.**
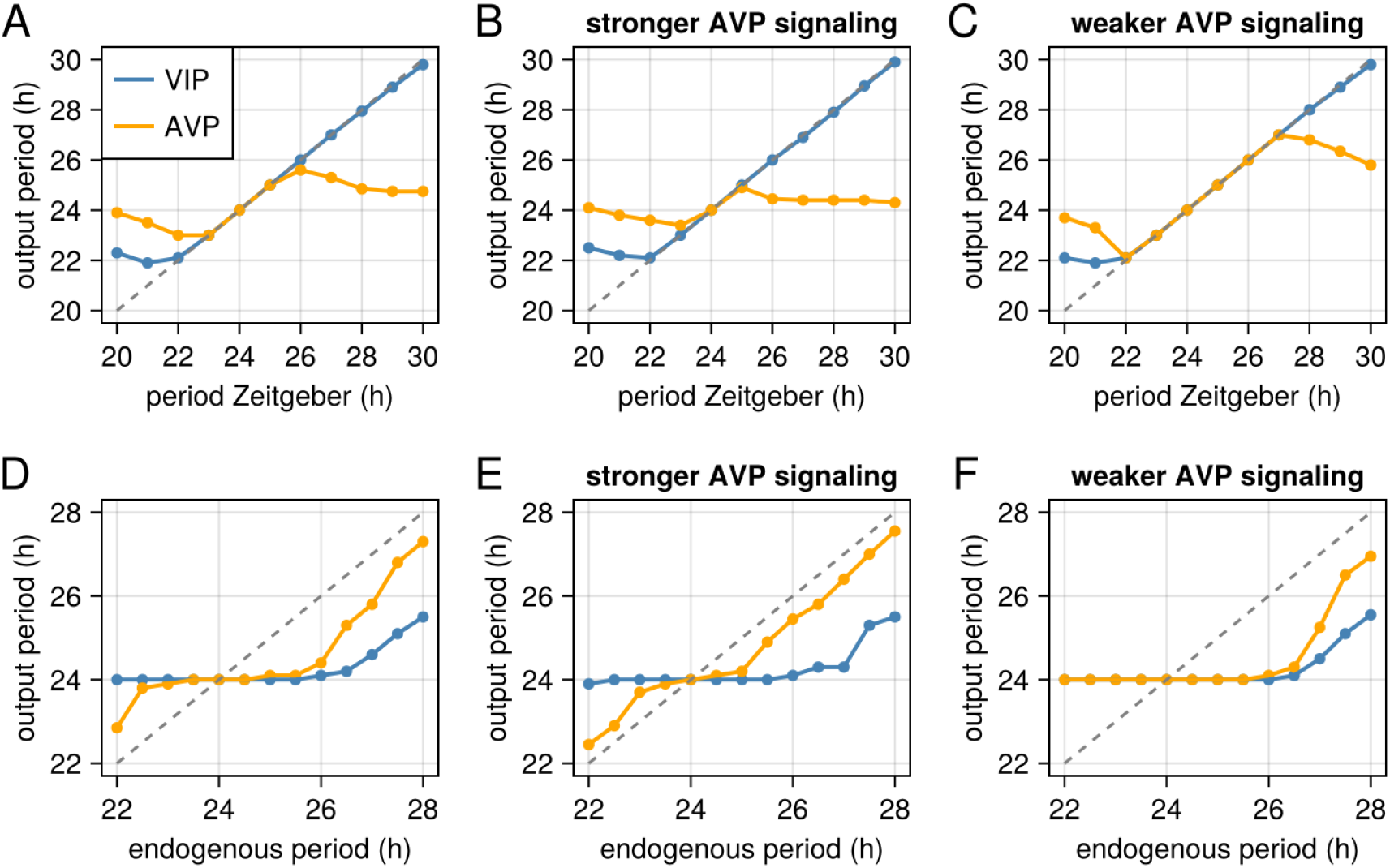
AVP signaling strength modulates SCN synchronization in response to changes in the zeitgeber period or endogenous period. This figure illustrates the relationship between the period of the zeitgeber (external time cue), the endogenous period (tau), and the synchronization of VIP and AVP neuronal outputs, highlighting the role of AVP signaling strength. A) VIP and AVP neurons typically synchronize to zeitgeber periods close to 24 hours. B) Stronger AVP signaling (double the standard value of κ_AA_ and κ_Av_) narrows the range of synchronizing zeitgeber periods, C) while weaker AVP signaling (half the standard value of κ_AA_ and κ_Av_) broadens this range. D) Similar effects are observed when varying tau. E) stronger AVP signaling leads to a shorter window of synchronization, while F) compared to weaker AVP signaling.

Importantly, the model predicts that the strength of AVP signaling modulates the range of zeitgeber periods over which VIP and AVP neurons remain synchronized. Strengthening AVP signaling narrows the synchronization range (Figure 6B), increasing the susceptibility to desynchronization. Conversely, weakening AVP signaling, such as through V1aR antagonism, broadens this synchronization range (Figure 6C). Similar behavior is observed when simulating the SCN output for different values of endogenous period (tau). For values of tau close to 24 hours, the SCN output (VIP and AVP) is synchronized (Figure 6D). However, increasing the strength of AVP signaling leads to a shorter window of synchronization (Figure 6E), while weakening it leads to a broader range of synchronization (Figure 6F). These findings highlight the crucial role of AVP signaling in regulating SCN synchrony and suggest that modulating AVP signaling strength, such as through V1aR antagonism, could be a strategy to restore SCN synchrony.

### Relationship between endogenous period, circadian phase and V1aR antagonism

To investigate the interplay between endogenous period (tau), circadian phase, and V1aR antagonism, we conducted simulations with varying tau lengths. Our model predicts that deviations of tau from 24 hours influence the phase of circadian biomarkers (melatonin and core body temperature). Specifically, increasing tau leads to a phase delay, with an approximate 30-minute increase in tau resulting in an approximate 2-hour phase delay in the circadian rhythms (Figure 7A-D). Conversely, decreasing tau leads to a phase advance. Importantly, the model predicts that V1aR antagonism induces phase shifts, with the magnitude and direction dependent on the deviation of tau from 24 hours (Figure 7E,F). When tau exceeds 24 hours, V1aR antagonism causes a phase advance, with larger deviations resulting in greater advances. Conversely, when tau is less than 24 hours, V1aR antagonism induces a phase delay, with larger deviations resulting in greater delays. Interestingly, we find that this effect is dependent on the inhibition AVP synchronization, and independent on the effect of V1aR antagonism on AVP to VIP signaling (Figure S4). Taken this together, it suggests that V1aR antagonism may be particularly effective in phase advancing individuals with long tau and phase delaying individuals with short tau by weakening the synchronization of AVP cells.

**Figure 7.**
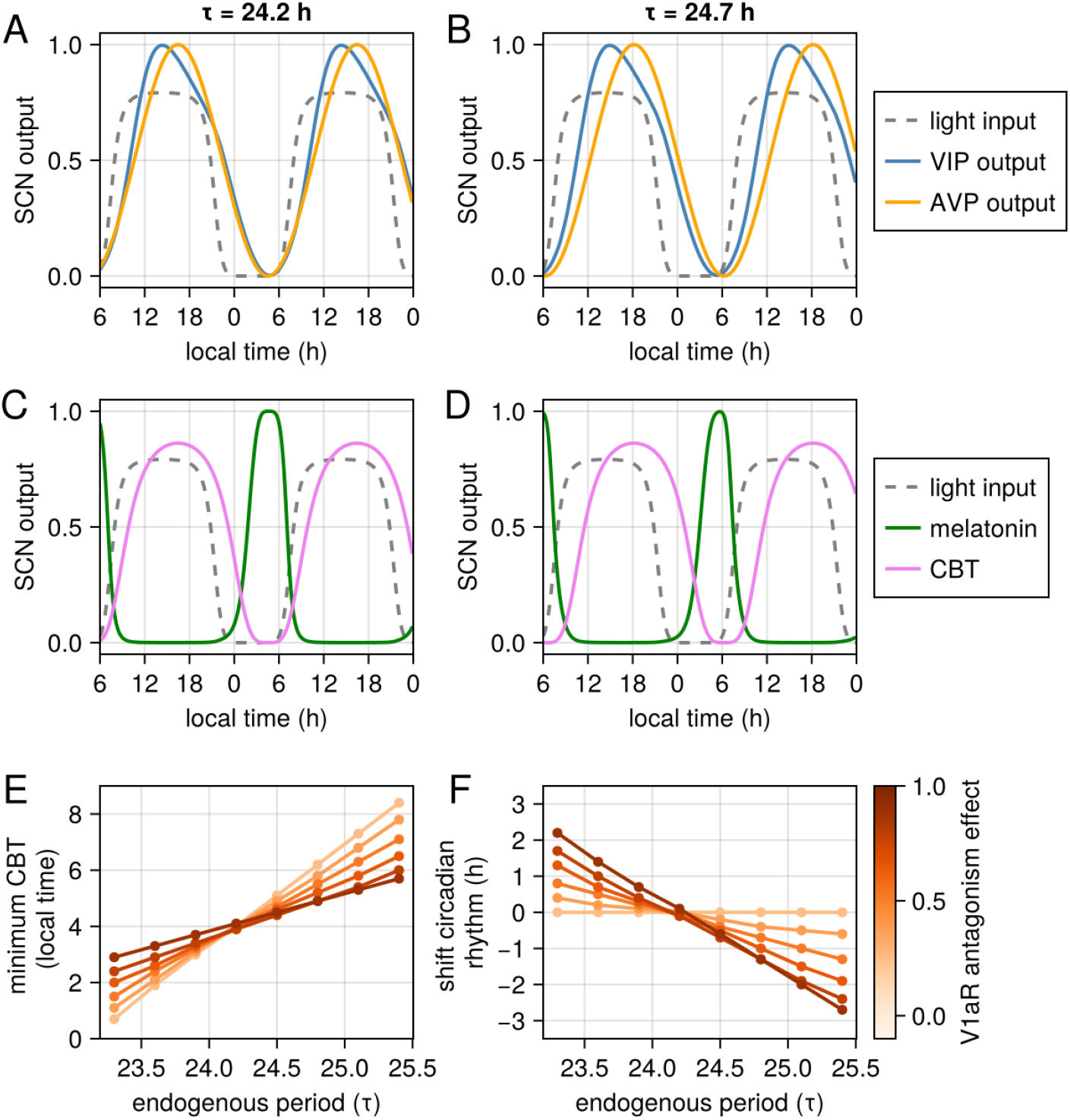
Relationship between the endogenous period (tau) and the timing of the minimum of core body temperature (CBT) and the effect of V1aR antagonism. (A-D) An increase in tau in 30 min leads to a phase advance of approximately 2h of circadian rhythm. E-F) V1aR antagonism induces phase shifts in the CBT rhythm, with the direction (advance or delay) and magnitude determined by the deviation of tau from 24 hours. tau > 24 hours leads to phase advances, with larger deviations resulting in greater advances. Conversely, tau < 24 hours leads to phase delays, with larger deviations resulting in greater delays.

## Discussion

This study shows that a single dose of the V1aR antagonist balovaptan is sufficient to significantly accelerate the resynchronization of behavioral rhythms following an experimental 6-hour jet lag in rodents. While simultaneous inhibition of V1a and V1b vasopressin receptors has been shown to speed up recovery from an experimental jet lag condition (Yamagushi et al, 2013), our finding that a single dose of balovaptan is sufficient to induce faster circadian resynchronization is in line with more recent mechanistic studies. Specifically, Yamaguchi et al. (2023) showed that the robustness of the circadian clock is mediated by two independent AVP pathways: one intra-SCN (V1a) and the other extra-SCN (V1b). These pathways were shown to contribute independently and additively to the speed of re-entrainment after a jet lag paradigm. SCN-specific V1a knockout mice showed significantly faster re-entrainment compared to wild-type mice, confirming the direct role of the intra-SCN V1a pathway (Yamagushi et al, 2013). This demonstration that both intra-SCN (V1a) and extra-SCN (V1b) pathways contribute additively to this effect further supports the therapeutic potential of targeting the V1a pathway alone. In addition, we introduced a mechanistic mathematical model of the SCN that simulates the regulation of key circadian biomarkers and the effects of V1aR antagonism. To validate our model, we showed that the model was able to replicate many well-established behaviors of the SCN, including the disruption of synchrony between AVP and VIP neurons under forced synchronization experiments, constant light exposure, and jet lag conditions. Once we were confident that our model was able to capture essential features of the SCN and V1aR antagonism effect on resynchronization after jet lag, we used the model to make predictions regarding the effect of V1aR antagonism in humans.

The first insight from our model is that variations in the strength of AVP signaling can influence the SCN’s ability to maintain synchronization. Our model predicts that stronger AVP signaling increases the susceptibility of the SCN to desynchronization (Figure 6). This suggests that AVP signaling strength could be a factor leading to SCN desynchronization, contributing to circadian dysfunctions related to circadian disruption, and could be the subject of further investigation. Additionally, the model indicates that weakening AVP signaling, achievable through V1aR antagonism, could enhance the SCN’s robustness against internal desynchrony and serve as a therapeutic target for disorders linked with SCN desynchrony. Many disorders, such as non-24-hour sleep-wake rhythm disorder (N24SWRD), have been associated with circadian disruption. N24SWRD has been linked to a longer endogenous period (Micic et al., 2013), which could be one potential cause of SCN desynchronization. Our model suggests that an endogenous period far from 24 hours could lead to SCN internal desynchrony and that weakening vasopressin signaling could restore this synchrony (Figure 6). Bipolar disorder (BD) has also been associated with SCN desynchrony, where individuals with BD often exhibit irregular sleep patterns and mood fluctuations corresponding with circadian rhythm disruptions (Harvey, 2008; McClung, 2007). Genetic studies implicate circadian genes, such as CLOCK and ARNTL, in BD pathophysiology, reinforcing the link between circadian disruption and mood disorders (Waddington Lamont et al., 2007). Despite being speculative, our model suggests that V1aR antagonism could be beneficial for these disorders.

The second insight from our model concerns the relationship between the endogenous period, phase of circadian rhythm, and V1aR antagonism. Our model predicts that an increase in the endogenous period results in a phase delay in circadian rhythms, where a 30-minute increase in tau leads to an approximate 2-hour phase delay. This observation is consistent with findings in individuals with Delayed Sleep-Wake Phase Disorder (DSWPD) (Micic et al., 2013; Micic et al., 2021). Furthermore, the model predicts that V1aR antagonism can induce a phase advance in individuals with longer endogenous periods, with the phase advance’s magnitude proportional to the length of the endogenous period. Experimental observations have suggested a longer tau as one potential cause of circadian delay in DSWPD (Micic et al., 2013; Micic et al., 2021). Although there is debate over whether tau is the cause of DSWPD or if it causes phase advances in only a subset of DSWPD individuals, our model suggests that targeting the V1aR pathway could present a novel therapeutic strategy for treating DSWPD by phase advancing their circadian rhythm.

Our model has several limitations. While we attempted to represent the key features of SCN biology, it is based on a simplified view of the SCN. Specifically, we assumed it consists of two regions: the vlSCN (core), primarily containing VIP neurons, and the dmSCN (shell), primarily containing AVP neurons. This approach overlooks the presence of various other cell types and entrainment agents within the SCN. Additionally, our model integrates experimental data from both humans and rodents, presuming a shared underlying biology, which may not always hold true. The model also does not consider downstream effects of circadian rhythm, such as sleep, or the reciprocal influence of melatonin feedback on circadian rhythmicity. Despite these limitations, our study provides a theoretical framework for evaluating the potential effects of targeting the SCN with pharmacological interventions. This framework can be readily adapted to include the pharmacodynamics of V1aR antagonism, assess its impact on SCN and circadian biomarkers, and support the design of future trials. The translation of these findings to humans could be tested by using a brain penetrant and selective V1a antagonist small molecule like balovaptan (Schnider et al, 2020), which has been shown to be safe and well tolerated in clinical studies (Bolognani et al, 2019; Jacob et al, 2022).

## Methods

### Mouse resynchronization studies

We used male young-adult (4 weeks old at the beginning of experiments) C57BL/6J mice. Animals were exposed to a light-dark cycle (LD) 12:12 (lights on from 8h to 20h geographical time; light intensity at 200lux; during dark period dim red light of 5lux) in individual cages with food (UAR, Epinay sur Orge, France) and water ad libitum and temperature (22 ± 1°C) and humidity (55% ± 5%) controlled. LD cycle was represented as zeitgeber times (ZT) where ZT0 represents the lights on and ZT12 lights off. Experiments were performed in accordance with the Principles of Laboratory Animal Care (National Institutes of Health publication 86-23, revised 1985) and the French national laws.

### Behavioural recording and analysis

The locomotor activity rhythms were measured using infra-red detectors on the top of the cages and/or running wheels. Data were acquired with Circadian Activity Monitoring System (CAMS, INSERM Lyon). Activity data were displayed as actograms and analyzed using the program ClockLab (Actimetrics, Evanston, IL, USA)

#### Re-entrainment (jet lag tests)

For 6 hours phase-shift (jet lag protocol) we quantified the number of days after the shift for total re-entrainment, considering both the onset and offset of activity. The onsets and offsets of activity were identified automatically using ClockLab software. Thus, full resynchronization to a new LD cycle, after 6-hour phase shifts of the time of lights off, was considered when activity onset took place at the new time of lights off ±30 minutes. We calculated the number of days (transitory cycles) required to re-synchronize each mouse. The speed of re-entrainment was evaluated by calculating the onset of activity and the advance of it, in hours, daily. To evaluate the effects of manipulation on locomotion, we quantified the locomotor activity changes from 0 to 6h after each treatment in 30 min intervals.

### Statistical analysis

Values in all experiments are means ± SEM. Statistical comparisons were performed using two-way analysis of variance (ANOVA) followed by post hoc Dunnett’s test for multiple comparison for the phase advance experiment. One way analysis of variance (ANOVA) was used for the numbers of days needed to reentrain. The critical value for statistical significance was p<0.05 for all experiments. We used GraphPad prism for the calculations (version 10.5).

## Mathematical Model

Mathematical models have been instrumental in deciphering the complex mechanisms governing SCN function and circadian timing. Early work established the role of negative feedback loops in gene regulation for generating sustained oscillations (Goodwin, 1965). Subsequent models expanded upon this foundation by incorporating light input and emphasizing the importance of intercellular coupling for achieving SCN neuron synchronization (Kronauer, 1990; Kuramoto, 1975).

This study utilizes a coupled oscillator model, where *N*_*V*_ and *N*_*A*_ represent the respective populations of VIP and AVP neurons. The model, based on the Kuramoto framework (Kuramoto, 1975), describes the temporal evolution of individual cell phases. Many previous studies have used Kuramoto based models to simulate synchronization in the SCN, which have been a source of inspiration of the model implemented here (Amdaoud et al, 2007; Rougemont and Naef 2008; Lu et al, 2016; Kori et al, 2017; Rohling and Meylahn, 2020). For a more in-depth theoretical analysis of Kuramoto based model that simulates the SCN divided into core and shell see Taylor et al, (2017) and Goltsev et al (2022).

The phase *θ*_*k*_ of each cell is translated into expression levels using the relationship 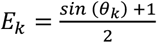, which varies from 0 to 1. The temporal dynamics of the phase are described by the equation:

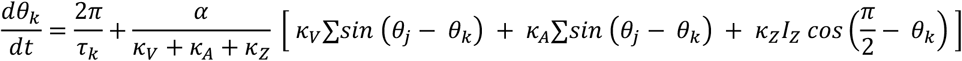

where *τ*_*k*_ is the endogenous period of oscillator *k, α* is the coupling constant, and *κ*_*V*_, *κ*_*A*_ and *κ*_*Z*_ represent the coupling strengths of VIP, AVP, and light input, respectively. For AVP cells, *κ*_*ZA*_ = 0 and representing the absence of light effect on the phase of AVPs.

Light input is the primary zeitgeber in the model, affecting the phase of the molecular clock in VIP cells. Light input is represented by the term *κ*_*Z*_*I*_*Z*_ where *κ*_*Z*_ is the strength of light input, and *I*_*Z*_ =*Z I*(*t*), where *Z* is the strength of light input and *I*(*t*) is the temporal regulation of the light input and assume values from 0 (dark/night) and 1 (light/day). Note that the light input is dependent on the term 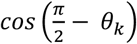. Mechanistically, this means that light input tends to drive the phase of the clock to 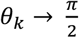 which represents the maximum expression of clock genes. Therefore, light input upregulates the expression of clock genes, and consequently *VIP* expression in VIP cells. Put another way, if the expression of *VIP* is increasing, a light input further accelerates its increase (therefore phase advancing). On the other hand, if the expression of *VIP* is decreasing, light delays its decrease (therefore phase delaying).

To visualize the dynamics of coupled oscillators it is convenient to define order parameters based on the following relationship:

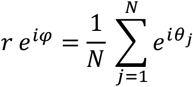

where, the radius *r*(t) measures the phase coherence, and *φ*(*t*) is the average phase. Based on that, and following the analysis in (Strogatz, 2000), we can rewrite the model in a more computationally efficient version:

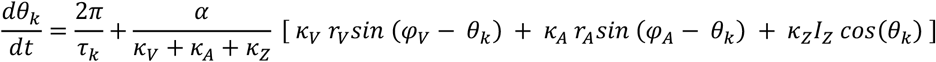

Where *r*_*V*_ and *φ*_*V*_ are the phase coherence and average phase of VIPs, and *r*_*A*_ and *φ*_*A*_ are the phase coherence and average phase of AVPs. The output of VIPs and AVPs can be defined by 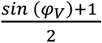 and 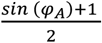, respectively. We also used the following equality: 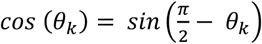.

The model relies on the following key assumptions: no spatial diffusion of VIP and AVP, no spatial localization of neurons (all neurons in the same population are equally affected), heterogeneity in the period of the molecular clock as a source of variability, the key circadian biomarkers are independently affected by the output of VIPs and AVPs, and unless indicated otherwise the function *I*(*t*) represents a modern human light schedule of 16 h of light and 8h of dark (as shown in Figure 3B).

### Effect of V1aR antagonists

We assumed that V1aR antagonists suppress the communication via AVP by blocking its receptor. Mathematically, we defined a variable V1aRa that represents the effect of V1aR antagonists and assume values from 0 (no effect) to 1 (full receptor blockage). Under the presence of V1aR antagonists we assumed that *κ*_*AA*_ → *κ*_*AA*_ V1aRa and *κ*_*AV*_ → *κ*_*AV*_ V1aRa.

### Standard parameter values

Unless indicated otherwise, the following values of the parameters were used in the simulations:

| Parameter | Value | Unit |
| --- | --- | --- |
| $N_V, N_A$ | 100, 200 | — |
| $\tau_k$ | 24.2 +- 0.5 | h |
| $\alpha$ | 0.25 | 1/h |
| $Z$ | 0.8 | — |
| $\kappa_{VV}, \kappa_{VA}$ | 1.0, 0.5 | — |
| $\kappa_{AV}, \kappa_{AA}$ | 0.5, 5.0 | — |
| $\kappa_{ZV}, \kappa_{ZA}$ | 25.0, 0.0 | — |

### Software

Figure generation and statistical analysis were performed in Julia (version 1.11.0) (Bezanson et al, 2017)

### Source Code

The source code can be found at https://github.com/mboareto/model_SCN

## Supplementary material

**Figure S1:**
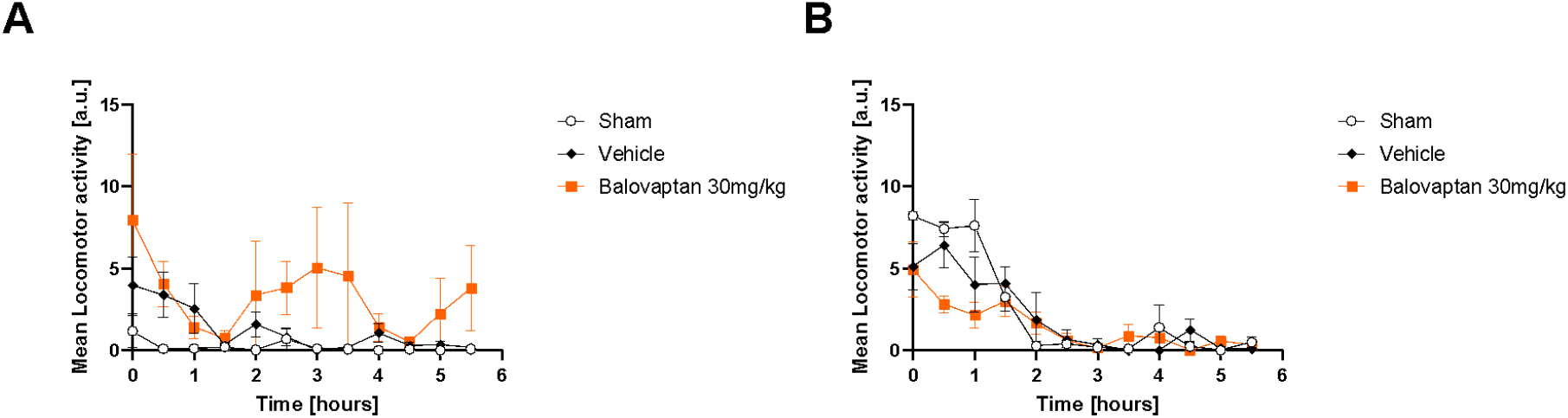
locomotor activity of mice in bins of 30 minutes (Mean and SEM) (A) first experiment (B) second experiment.

**Figure S2:**
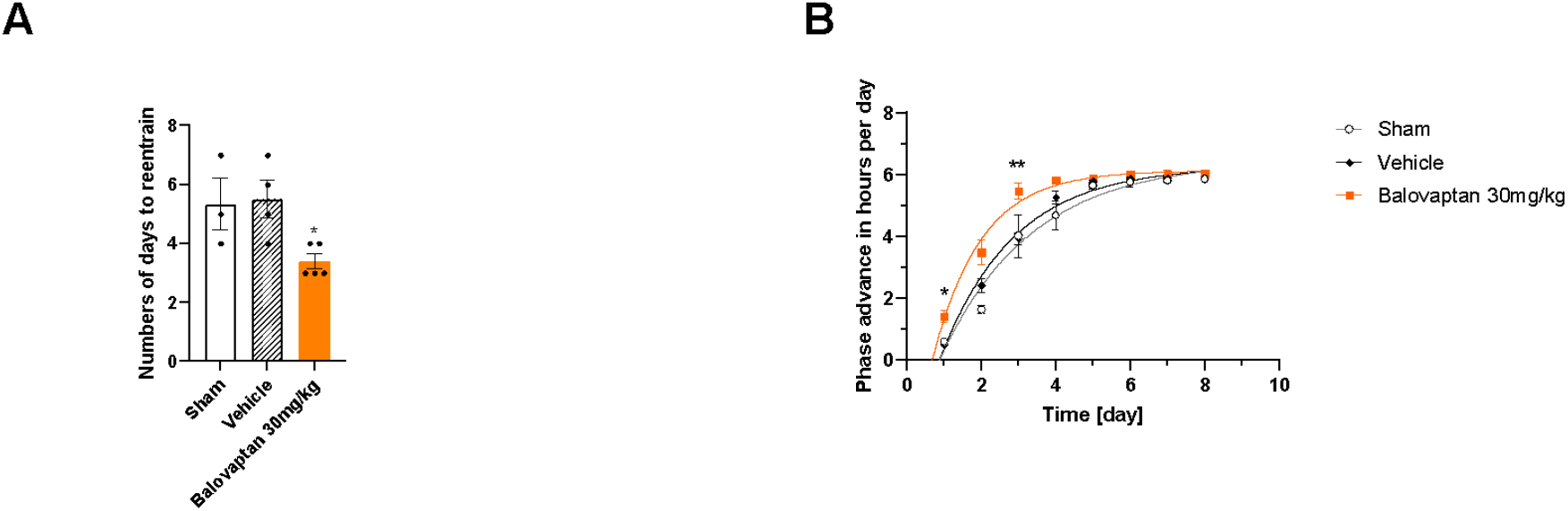
A) Rate of re-entrainment (number of days) to show full synchronization to the new LD cycle (Dunnett’s post-hoc test vs vehicle.* p< 0.05). B) The speed of entrainment was evaluated daily and on days 1 and 3 the 30 mg/kg group showed a significantly faster re-entrainment than the vehicle group (Dunnett’s post-hoc test vs vehicle.* p< 0.05, ** p< 0.01). (sham n = 3, vehicle n = 4, 30 mg/kg n = 5).

**Figure S3.**
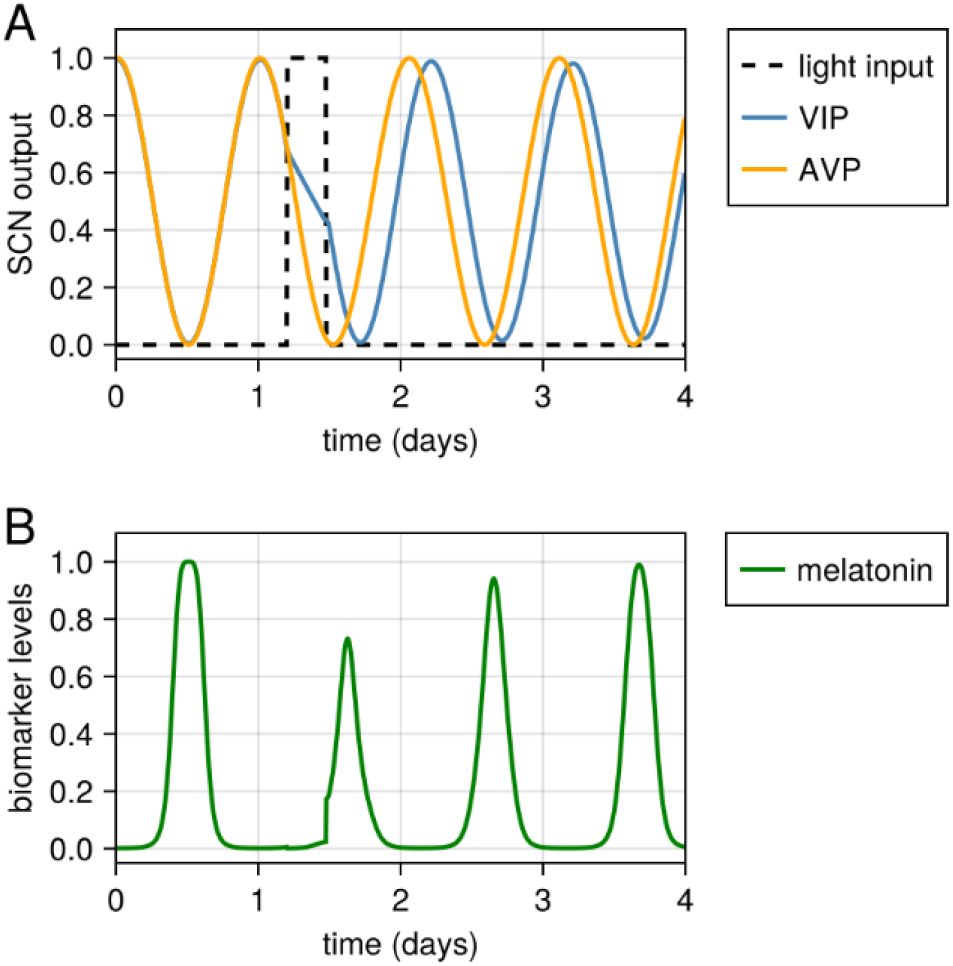
Impact of a light pulse on SCN output and melatonin levels. This figure shows the effect of a light pulse administered before the expected melatonin rise on A) SCN output and B) melatonin levels. The model accurately replicates the suppression of melatonin onset and the resulting in a slightly shorter consequent melatonin peak, aligning with experimental findings (Khalsa et al., 2003).

**Figure S4.**
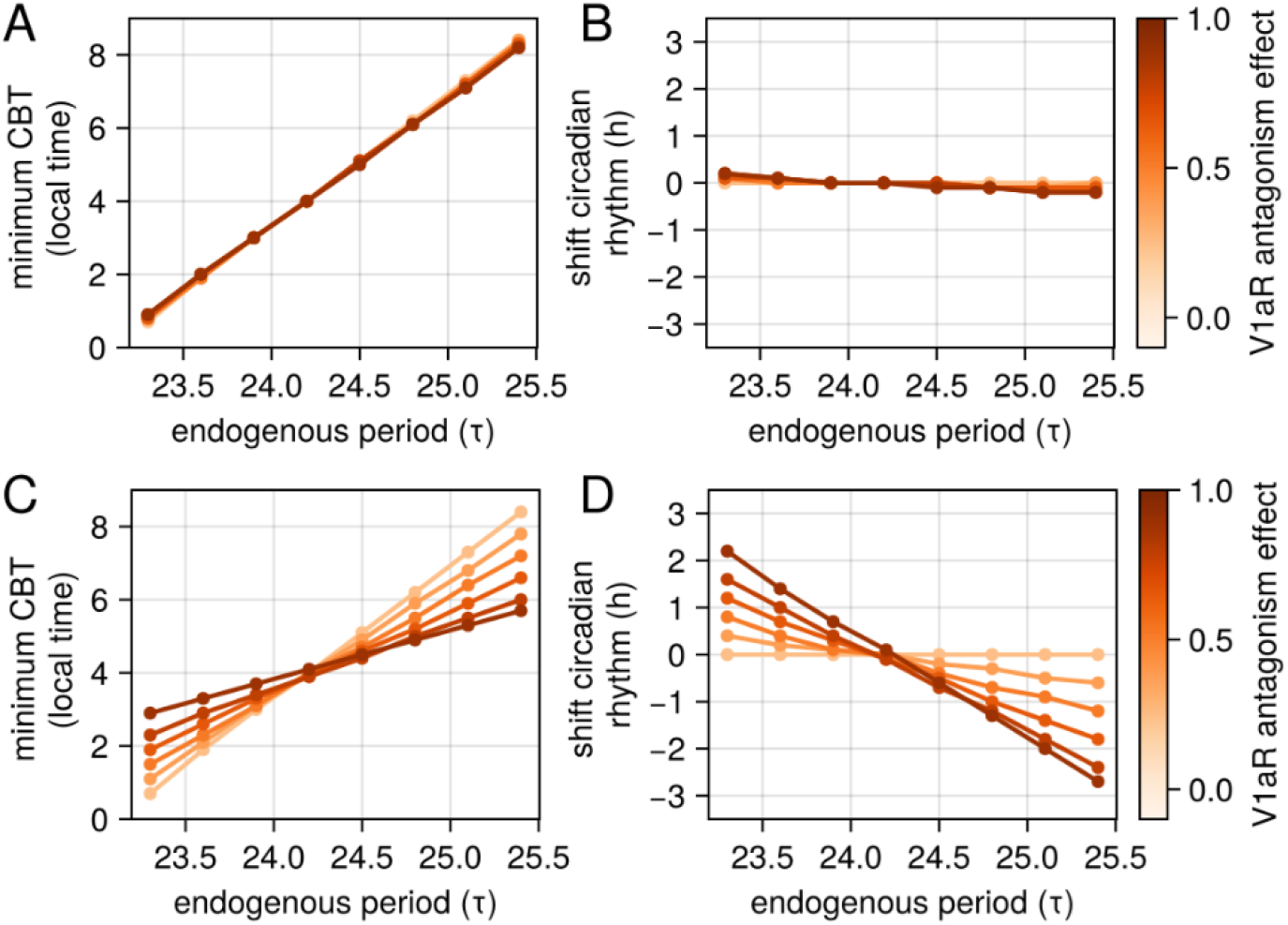
A,B) same as figure 7E,F with V1aR antagonism effect only acting on κ_AV_. C,D) same as figure 7E,F with V1aR effect only on κ_AA_.

